# Context-dependent effects of whole-genome duplication during mammary tumor recurrence

**DOI:** 10.1101/2021.05.03.442527

**Authors:** Rachel Newcomb, Emily Dean, James V. Alvarez

## Abstract

Whole-genome duplication (WGD) generates polyploid cells possessing more than two copies of the genome and is among the most common genetic abnormalities in cancer. The frequency of WGD increases in advanced and metastatic tumors, and WGD is associated with poor prognosis in diverse tumor types, suggesting a functional role for polyploidy in tumor progression. Experimental evidence suggests that polyploidy has both tumor-promoting and suppressing effects, but how polyploidy regulates tumor progression remains unclear. Using a genetically engineered mouse model of Her2-driven breast cancer, we explored the prevalence and consequences of whole-genome duplication during tumor growth and recurrence. While primary tumors in this model are invariably diploid, nearly 40% of recurrent tumors undergo WGD. WGD in recurrent tumors was associated with increased chromosomal instability, decreased proliferation and increased survival in stress conditions. The effects of WGD on tumor growth were dependent on tumor stage. Surprisingly, in recurrent tumor cells WGD slowed tumor formation, growth rate and opposed the process of recurrence, while WGD promoted the growth of primary tumors. These findings highlight the importance of identifying conditions that promote the growth of polyploid tumors, including the cooperating genetic mutations that allow cells to overcome the barriers to WGD tumor cell growth and proliferation.

## Introduction

Whole-genome duplication (WGD) events give rise to cancer cells possessing more than two copies of the genome. WGD is estimated as one of the most common genetic events in cancer [1]. Genomic analysis of human tumor sequencing data has revealed that this phenomenon is highly prevalent across cancers of different tissue types and with diverse oncogenic driving mutations, including breast cancer. Previous studies have estimated that up to 45% of breast cancers undergo at least one whole-genome duplication event [1]. As in the case of many cancers [2, 3], WGD is associated with aggressive disease and poor prognosis in breast cancer [2, 4–6]. WGD is also prevalent in metastatic disease [7] and in breast cancer has been observed in a greater proportion of metastases than primary tumors [5]. The prevalence of WGD in cancer, and its association with poor patient outcomes, underscores the need for better understanding how duplication of the cancer genome alters the biology of cancer cells to promote tumor progression.

The mechanistic basis by which WGD influences tumor evolution are beginning to be deciphered. Experimental evidence suggests polyploidy and near-tetraploid karyotypes can result in both tumor promoting and suppressing effects. Polyploidy is thought to promote tumorigenesis by increasing genomic instability and acting as a precursor to aneuploidy [8–10]. On the other hand, polyploidy and aneuploidy are known to activate strong cellular stresses including the p53 and Hippo pathways, as well as immune surveillance mechanisms [11–13]. Thus, the effects of WGD on cancer cells, and by extension on tumor evolution, are likely shaped by the context in which these events occur.

Understanding these context-dependent effects of WGD would benefit from experimental models that mimic human cancers in undergoing WGD during tumor progression. We have previously used a genetically engineered mouse model that recapitulates many aspects of breast cancer recurrence as it occurs in women [14–18]. In the current study, we show that recurrent breast tumors in these models spontaneously undergo WGD. Through a series of in vivo and in vitro experiments, we explore the cellular effects of WGD and its consequences on tumor growth and recurrence.

## Results

### Tumor recurrence is associated with alterations in tumor cell ploidy

To understand mechanisms of tumor progression, we employed a genetically engineered mouse model with inducible Her2 expression. In this model, MTB;TAN mice harbor two transgenic alleles: an MTB allele which expresses a reverse tetracycline transactivator (rtTA) under the control of the mammary gland-specific MMTV promoter; and a TAN allele which expresses the oncogene Her2 under the control of a tetracycline-responsive promoter. In the presence of tetracycline (or its analog doxycycline, dox), rtTA drives expression of Her2 in the mammary gland. This inducible expression system can be used to model the cascade of primary tumor formation, response to therapy and recurrence experienced in the clinic. Administration of dox leads to Her2 expression and the formation of invasive mammary adenocarcinomas. Subsequent withdrawal of doxycycline induces Her2 downregulation and complete tumor regression, which may mimic the actions of targeted therapies against Her2. However, residual cells survive oncogene down regulation and remain in a non-proliferative state in the mammary gland. These cells eventually reinitiate proliferation to form a recurrent tumor and these recurrent tumors grow independently of Her2 expression [17].

Given the prevalence of WGD events in human breast cancers, we hypothesized that WGD events may also occur during tumor development in this model, providing an opportunity to examine the effects of polyploidy at different stages of tumor progression. To address this, we generated a cohort of primary and recurrent tumors to examine tumor cell ploidy. Primary Her2-driven tumors were generated by administering doxycycline to MTB;TAN mice. One cohort of mice was sacrificed with primary tumors (Her2 on, +dox). Doxycycline was removed from a second cohort of mice with primary tumors to induce Her2 downregulation and tumor regression. These mice were palpated until the appearance of Her2-independent recurrent tumors. Primary or recurrent tumors were then harvested and digested to generate single cells. Flow cytometry and DNA content staining was performed, and the percentage of diploid and tetraploid cells was estimated using cell cycle modeling. This analysis demonstrated that all primary tumors (n=9) possessed near-diploid DNA content (Figure 1A-C). In contrast, over one-third of recurrent tumors (n=13) were constituted by cells with a near tetraploid DNA content, evidence of a whole-genome duplication event (Figure 1A-C).

**Figure 1.**
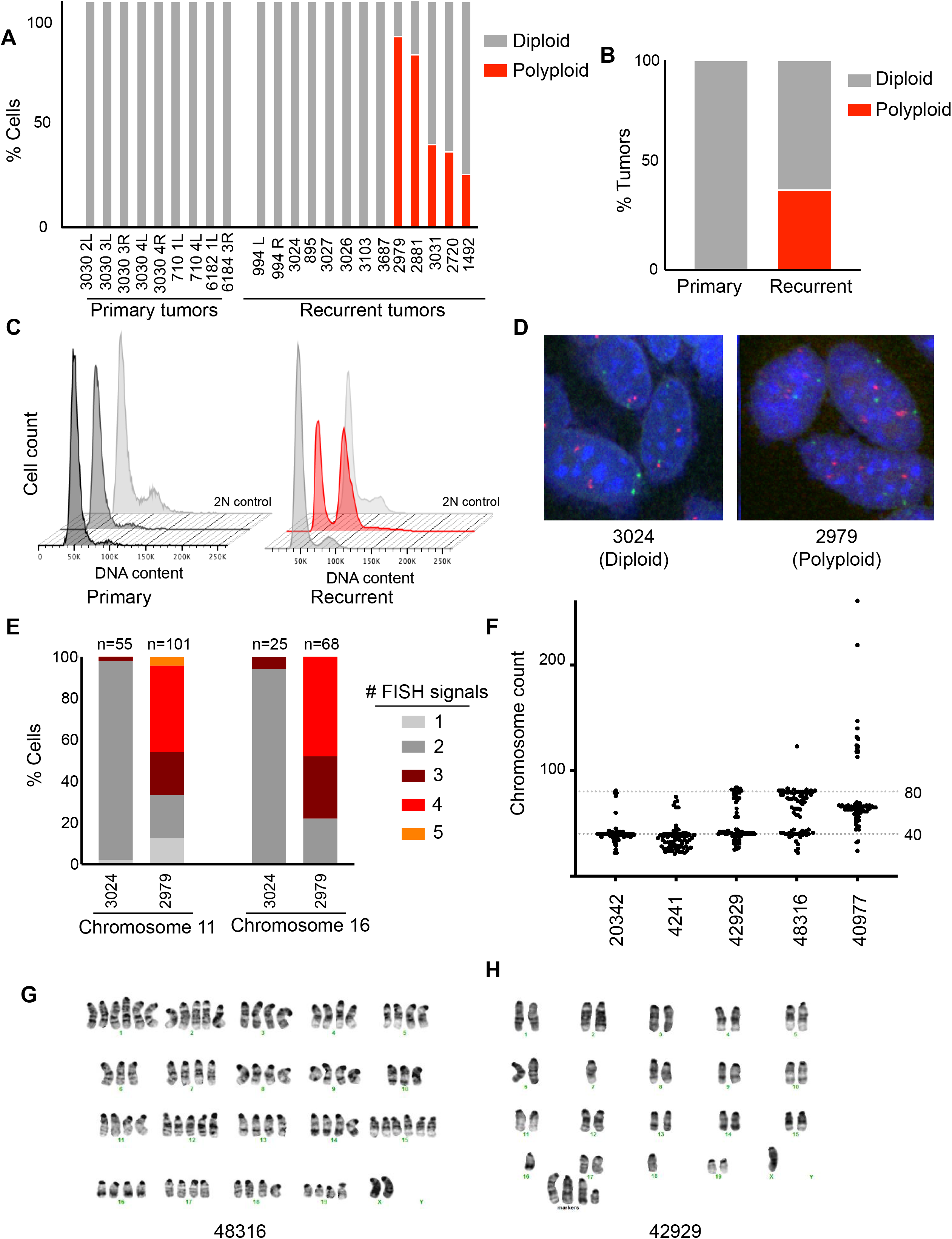
Tumor recurrence is associated with alterations in tumor cell ploidy. A) Quantification of DNA content staining to determine the ploidy of primary (n=9) and recurrent (n=13) tumors derived from MTB;TAN mice. The percentage of near-diploid and near-tetraploid cells in each tumor was estimated using cell cycle modeling. B) The frequency of whole-genome duplication in primary and recurrent tumors. C) DNA content staining of representative primary and recurrent tumors. A diploid cell line is shown as a control. D) Representative FISH images for chromosome 2 (red) and 11 (green) on a diploid and polyploid tumor. E) FISH using probes targeting chromosome 2 and 16 on a diploid tumor and a polyploid tumor. The number of FISH signals per cell was scored. F) Chromosome counts from metaphase spreads of cells cultured from recurrent tumors arising in MTB;TAN mice. G) and H) Representative karyotypes of cells cultured from recurrent tumors.

We next sought to validate our observation of WGD using a complementary method. Fluorescence in situ hybridization (FISH) was performed on digested tumor cells from a diploid tumor and one showing evidence of WGD, using probes against chromosomes 2 and 11 and chromosomes X and 16 (Figure 1D, E and Supplemental Figure 1A). Tumors that showed evidence of WGD by DNA content analysis had elevated chromosome counts for chromosomes 11 and 16 (Figure 1E). Elevated chromosome counts were observed in single nuclei (Figure 1D), indicating that recurrent tumors with WGD were comprised of mononucleate tetraploid cells. Importantly, similar results were obtained using probes targeting chromosome 16 and the X chromosome (Supplemental Figure 1A).

Both DNA content staining and FISH analysis showed that recurrent tumors contained both diploid and tetraploid cells. However, these tumors consist of tumor cells as well as non-tumor cells, including immune cells, fibroblasts, and endothelial cells. In fact, CD45+ cells can constitute up to 50% of all cells in recurrent tumors arising in MTB; TAN mice [15]. To determine if we were underestimating the fraction of polyploid cells in recurrent tumors, we depleted CD45+ immune cells from tumors, and performed DNA content analysis alongside non-depleted controls. CD45+ cell depletion increased the proportion of tetraploid cells in tumors that showed evidence of WGD but did not alter the proportion of tetraploid cells in diploid tumors (Supplemental Figure 1B). This suggests that the percentage of polyploid cells in recurrent tumors is likely higher than estimated from DNA content staining of bulk cells, and in some cases polyploid cells constitute ~90% of tumor cells in recurrent tumors (Supplemental Figure 1B).

To gain further insight into the chromosome content of recurrent tumor cells, we generated early-passage cell lines from a panel of recurrent tumors and performed metaphase spreads on these recurrent tumor cell cultures. Consistent with DNA content staining, these recurrent tumor cell lines exhibited a range of chromosome numbers. Two lines (20342 and 4241) had near-diploid chromosome counts (Figure 1F and Supplemental Figure 1C), while two lines (42929 and 48316) comprised cells with both diploid and tetraploid complements of chromosomes (Figure 1F and Supplemental Figure 1C). A final line, 40977, had a triploid chromosome count (Figure 1F and Supplemental Figure 1C). Karyotyping of 42929 and 48316 cells confirmed that these lines contained a mixture of diploid and tetraploid cells (Figure 1G and H).

We next explored whether WGD was correlated with features of the recurrent tumors or the primary tumors from which they arose. We found no significant difference between diploid and polyploid tumors in terms of maximum primary tumor volume, nor in recurrent tumor volume at time of sacrifice (Supplemental Figure 1E and F; Welch’s t-test, p=0.406 for primary tumors; p=0.97 for recurrent tumors). Retrospective analysis of tumors binned as polyploid or diploid showed no difference in time to tumor recurrence (Supplemental Figure 1 D and G; log-rank test, p=0.606; Hazards Ratio=0.66).

Together, these results demonstrate that a subset of recurrent tumors are composed of tetraploid tumor cells, indicating that a whole-genome duplication event occurs during the process of tumor recurrence. These findings suggest that whole-genome duplication may contribute to tumor recurrence.

### Generation of matched tumor cell models to investigate the functional consequences of polyploidy

To investigate the impact of WGD on recurrent tumor cells, we generated two models of matched diploid and tetraploid tumor cells, a spontaneous model (Figure 2A) and an induced model (Figure 2B). The recurrent tumor cell line 42929, which was composed of a mixture of diploid and spontaneously formed tetraploid cells (see Figure 1F and Supplemental 1A), was sorted into pure populations of diploid and tetraploid cells based on DNA content using live-cell DNA staining and fluorescence-activated cell sorting (FACS) (Supplemental Figure 2B). The ploidy of sorted populations of diploid and tetraploid 42929 cells remained relatively stable over time (Figure 2C-F). In the sorted diploid population, some tetraploid cells accumulated during passaging (Figure 2C), though cell cycle modeling estimated that greater than 75% of this sorted population was composed of diploid cells after 8 population doublings (Figure 2D). The ploidy of the sorted tetraploid population remained stable, with nearly 100% tetraploid cells (Figure 2E and F). We refer to this pair of matched cell lines as a spontaneous model of tetraploidy, reflecting the fact that tetraploid cells in this line arose spontaneously during tumor recurrence.

**Figure 2.**
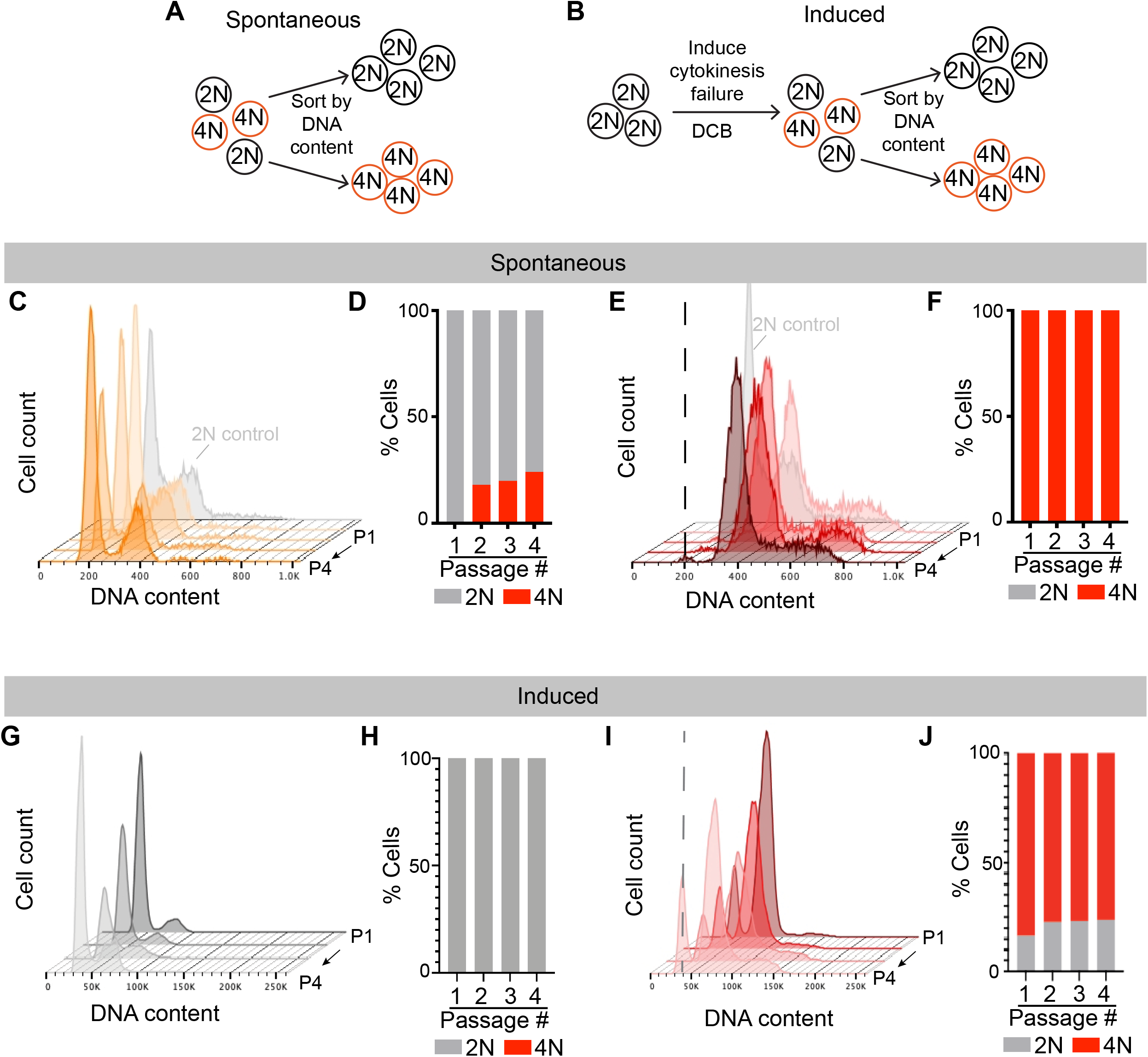
Creation of matched diploid and tetraploid recurrent tumor cell models. A) Schematic outlining the generation of matched diploid and tetraploid populations from a recurrent tumor with spontaneously occurring tetraploid cells. B) Schematic outlining the generation of isogenic diploid and tetraploid recurrent tumor cells following treatment of diploid tumor cell cultures with the cytokinesis inhibitor DCB. C) and D) Stability of tumor cell ploidy in sorted diploid cells from the spontaneous model over increasing passages. Representative DNA content staining is shown in (C), and the percentage of diploid and tetraploid cells at each passage in shown in (D). E) and F) Stability of tumor cell ploidy in sorted tetraploid cells from the spontaneous model over increasing passages. Representative DNA content staining is shown in (E), and the percent of diploid and tetraploid cells at each passage in shown in (F). Dotted line in (E) corresponds to the G1 peak of the 2N control. G and H) Stability of tumor cell ploidy in sorted diploid cells from the induced model over increasing passages. Representative DNA content staining is shown in (G), and the percentage of diploid and tetraploid cells at each passage is shown in (H). I and J) Stability of tumor cell ploidy in sorted tetraploid cells from the induced model over increasing passages. Representative DNA content staining is shown in (I), and the percent of diploid and tetraploid cells at each passage is shown in (J). Dotted line in (I) corresponds to the G1 peak of the 2N control.

A second model was created by inducing tetraploidy in the recurrent tumor cell line 4241, which was composed entirely of diploid tumor cells (see Figure 1F). The 4241 cells were treated with a low dose of the actin inhibitor dihydrocytochalasin B for 16 hours, resulting in cytokinesis failure and the induction of tetraploidy in ~15% of cells (Supplemental Figure 2C, D and E). Diploid and tetraploid cells from this population were then sorted based on DNA content using FACS (Supplemental Figure 2C and E). We again monitored the stability of DNA content in each sorted population with increased passaging (Figure 2G-J). The ploidy of the sorted diploid population remained stable, with nearly 100% of the cells having diploid DNA content (Figure 2G and H). The sorted tetraploid population accumulated a small fraction of cells with diploid DNA content, though greater than 70% of cells in this population were tetraploid after 8 population doublings (Figure 2I-J). We refer to this pair of matched cell lines as an induced model of tetraploidy.

### Polyploid recurrent tumor cells exhibit aneuploidy and elevated chromosomal instability

Equipped with these spontaneous and induced matched models of polyploidy, we next set out to characterize the consequences of polyploidy on recurrent tumor cells. Polyploidy is associated with aneuploidy in cancer and can promote chromosomal instability (CIN) [9, 19.] To determine whether polyploid recurrent tumor cells have elevated rates of aneuploidy, we performed metaphase spreads on polyploid and diploid cells from the spontaneous and induced model. As expected, the median number of chromosomes in metaphase cells from the polyploid population was approximately twice that of diploid cells from both models (Figure 3A-B; median chromosome count: spontaneous model, 74 vs 39; induced model, 68 vs 36). Interestingly, polyploid populations had a larger variance in their distribution of chromosome counts, indicative of a higher degree of aneuploidy in these populations (Figure 3A-B; F test of variances: spontaneous model, P=0.0073; induced model, P<0.0001). To assess rates of ongoing CIN in diploid and polyploid cells, we generated and expanded single-cell clones from diploid and polyploid populations and measured chromosome numbers in the expanded populations using metaphase spreads. Cells derived from diploid clones exhibited minimal variability in chromosome number from the diploid complement of 40 (Figure 3C-D). In contrast, cells derived from tetraploid clones exhibited a wide variance in chromosome counts. This was especially evident in the spontaneous model, where populations derived from three independent polyploid clones had a wide range of chromosome number (Figure 3C). Interestingly, one of the two polyploid clones in the induced model gave rise to largely diploid cells (Figure 3D, Polyploid clone 2), likely reflective of a depolyploidization event. Taken together, these results demonstrate that both spontaneous and induced tetraploid cells have higher rates of CIN and aneuploidy than their diploid counterparts, suggesting that CIN is a direct consequence of the tetraploid state.

**Figure 3.**
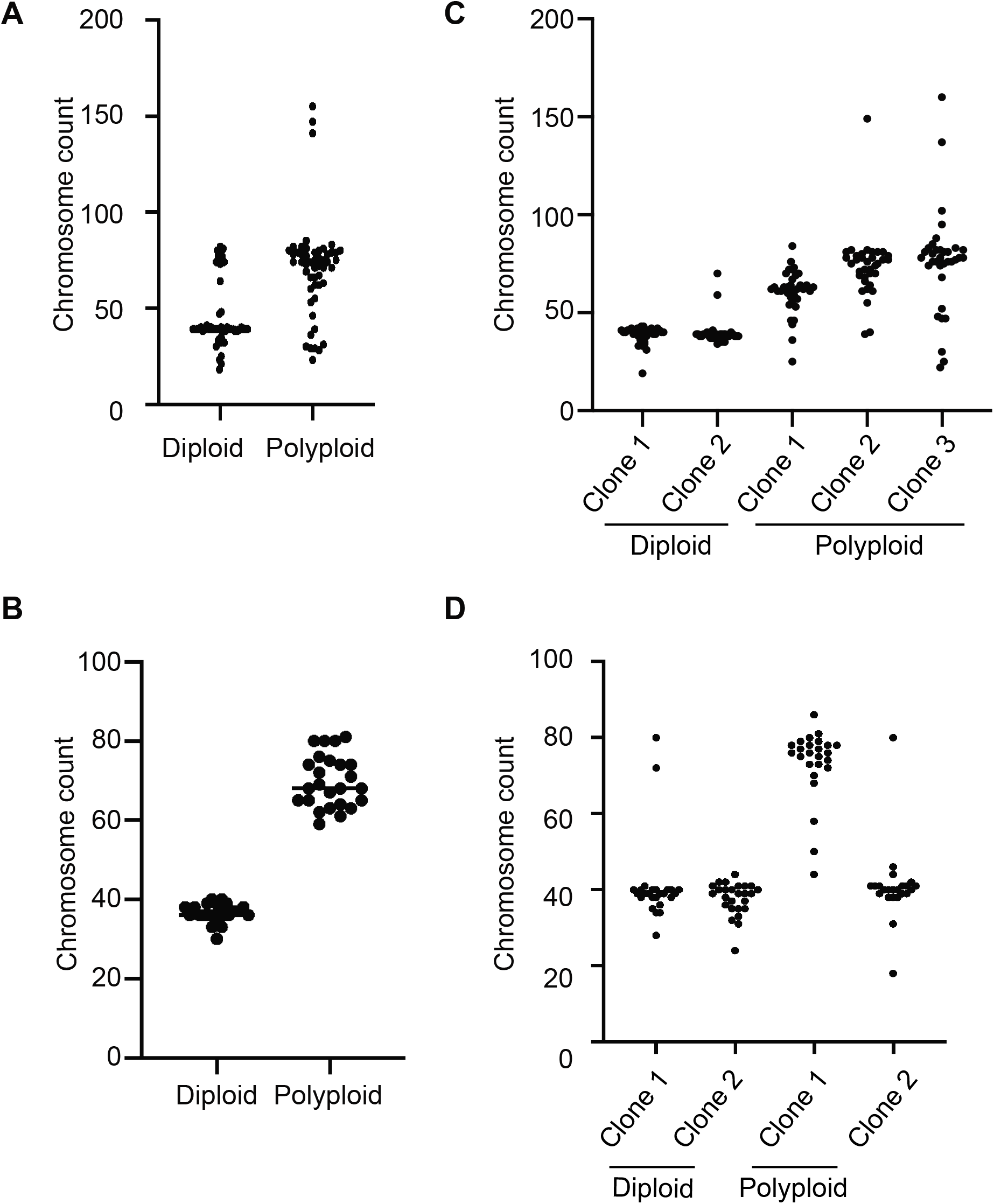
Tetraploid cells exhibit increased rates of chromosomal instability. A) Chromosome counts in sorted diploid and tetraploid cells from the spontaneous model. Bars indicate the median cell count. Differences in variance between diploid and tetraploid cells was determined using an F-test for variance, p = 0.0073. B) Chromosome counts in sorted diploid and tetraploid cells from the induced model. Bars indicate the median cell count. Differences in variance between diploid and tetraploid cells was determined using an F-test for variance, p < 0.0001. C) Chromosome counts in single-cell clones derived from diploid or tetraploid cells from the spontaneous model (n=35). D) Chromosome counts in single-cell clones derived from diploid or tetraploid cells from the induced model (n=25).

### Spontaneous and induced polyploidy slows tumor growth

Polyploid cells can either have enhanced or reduced tumorigenic capacity [8, 10, 13], and this may dependent upon the spectrum of oncogenic and tumor suppressor mutations present in the tumor. To gain insight into the tumorigenic potential of diploid and tetraploid recurrent tumor cells, we compared the growth of these cells in the mammary fat pad using an orthotopic tumor growth assay in athymic mice. Diploid and tetraploid recurrent tumor cells from the spontaneous or induced model were injected orthotopically into recipient mice, and tumor growth was measured biweekly using calipers. Surprisingly, polyploid cells from both the spontaneous and induced models took longer to form tumors compared to diploid cells (Figure 4A and D; log-rank test, p=0.0031 for spontaneous model; p<0.001 for induced model), and when polyploid tumors did form they grew more slowly than diploid tumors (Figure 4B, C, E and F; Welch’s t-test, p < 0.05). Thus, while polyploidy occurs spontaneously in a substantial fraction of recurrent tumors, polyploid recurrent tumor cells grow more slowly than their diploid counterparts.

**Figure 4.**
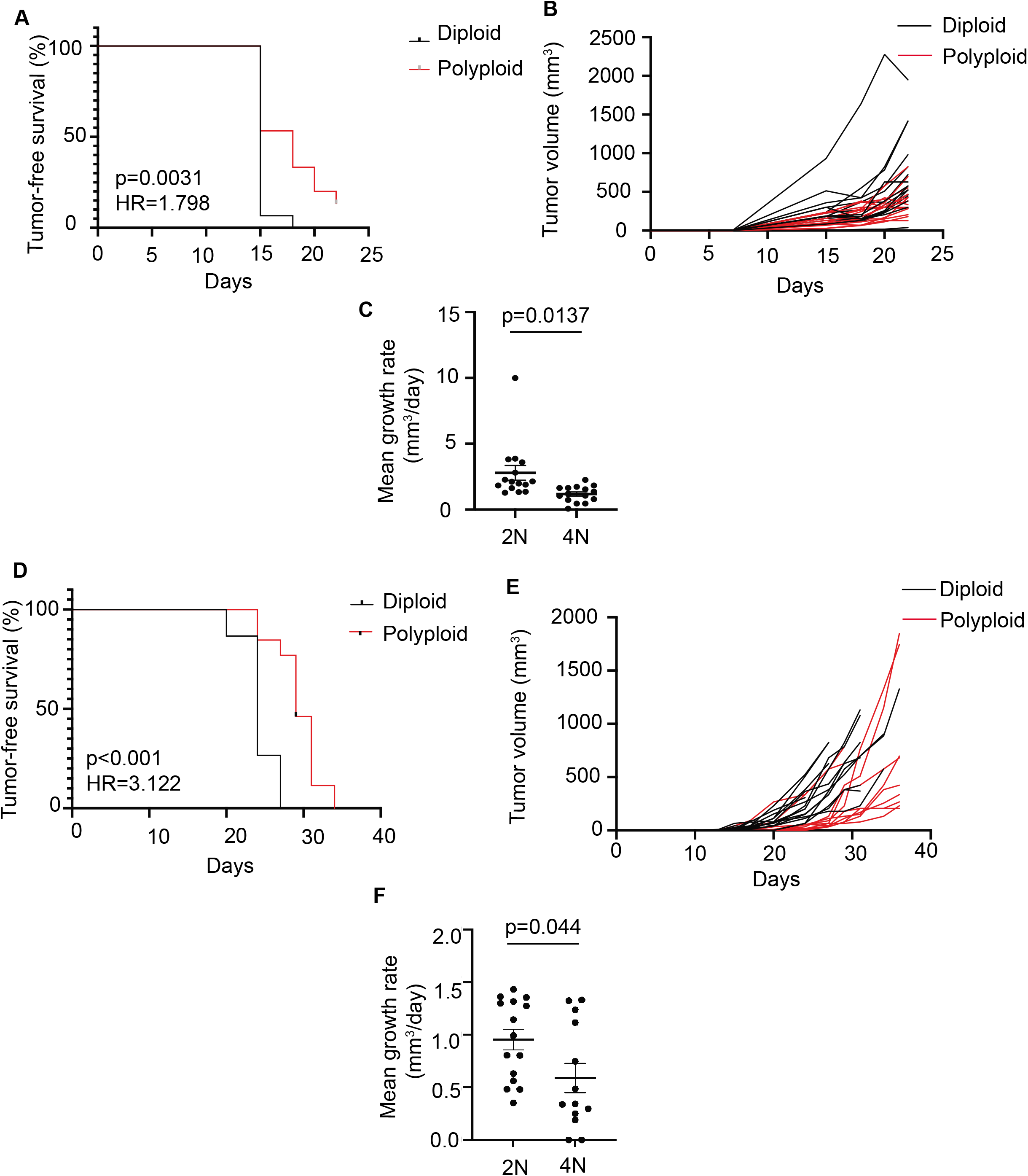
Polyploidy opposes tumor growth. A) Kaplan-Meier plot showing tumor-free survival for mice injected with diploid or polyploid tumor cells from the spontaneous model. Hazards Ratio (HR) and p-value were determined using a log-rank test. B) Tumor growth curves for diploid and polyploid tumors from the spontaneous model. C) Mean growth rate for diploid and polyploid tumors from the spontaneous model. Significance was determined using Welch’s t-test. D) Kaplan-Meier plot showing tumor-free survival for mice injected with diploid or polyploid tumor cells from the induced model. Hazards Ratio (HR) and p-value were determined using a log-rank test. E) Tumor growth curves for diploid and polyploid tumors from the induced model. F) Mean growth rate for diploid and polyploid tumors from the spontaneous model. Significance was determined using Welch’s t-test.

### Context-dependent effects of polyploidy on tumor cell proliferation and survival

We next examined the cellular basis of the effects of polyploidy on tumor growth. Both polyploidy and aneuploidy have been associated with reduced cell proliferation and cell cycle arrest [20], so we measured the proliferation rate of diploid and polyploid cells from the induced and spontaneous models. Induced and spontaneous tetraploid cells exhibited a proliferation defect relative to diploids (Figure 5A and Supplemental Figure 3A). Given the reported defects in cell cycle progression of polyploid cells, we examined the cell cycle distribution of diploid and polyploid cells in this model at varying serum concentrations to mimic reduced nutrient and growth factor availability. Quantification of cells in S-phase by BrdU staining and DNA content analysis by propidium iodide staining demonstrated that fewer tetraploids were in S-phase relative to diploids across various serum concentrations (Figure 5B and C; Two-way ANOVA with Sidak’s multiple comparisons test, p<0.05 for S-phase proportion at all serum concentrations; p<0.05 for BrdU+ at all serum concentrations). Consistent with this, we observed an increased proportion of tetraploids in G1 phase at all serum concentrations (Figure 5D; Two-way ANOVA with Sidak’s multiple comparisons test, p<0.05 at all serum concentrations).

**Figure 5.**
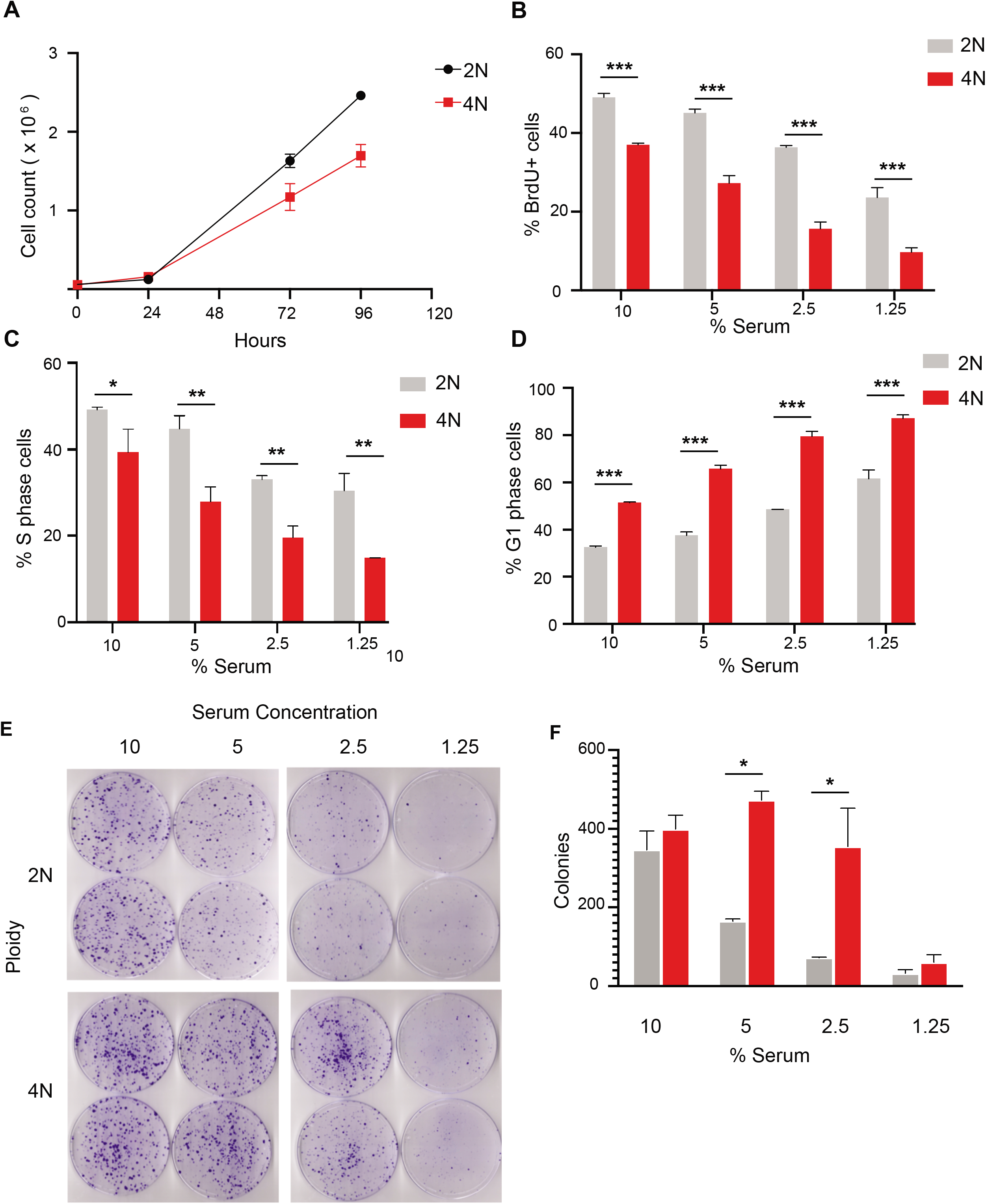
Tetraploid recurrent tumor cells demonstrate a competitive advantage in low serum culture. A) Cell count of diploid and induced tetraploid cells over time. B) The percentage of BrdU+ diploid and tetraploid cells at different serum concentrations. C) The percentage of diploid and tetraploid cells in S phase at different serum concentrations as measured by DNA content analysis and cell cycle modeling. D) The percentage of diploid and tetraploid cells in G1 phase at different serum concentrations as measured by DNA content analysis and cell cycle modeling. E) Colony formation assays of diploid and spontaneously formed tetraploid cells in different serum concentrations. F) Quantification of colony number from E. For B-E, significance was determined using two-way ANOVA with Sidak’s multiple comparison test. *p<0.05, **p<0.01, ***p<0.001

While tetraploidy can inhibit cell proliferation, it can also promote cell survival and adaptation to stress [21]. Furthermore, aneuploid cells that grow slowly in normal culture conditions have a selective advantage under conditions of stress [22]. We therefore used clonogenic assays to compare the long-term relative fitness of diploid and tetraploid cells in low serum conditions. Interestingly, in spite of their proliferation defect, tetraploid cells from the induced model formed more colonies in low serum conditions than diploids (Figure 5E and F; Two-way ANOVA with Sidak’s multiple comparisons test, p<0.05 at 5% and 2.5% serum). Similar results were observed in the spontaneous models (Supplemental Figure 3B and C; Student’s t-test, p=0.005). These results suggest that polyploidy is associated with increased survival under low nutrient/growth factor conditions, and this was independent of whether polyploidy evolved naturally or was experimentally induced.

### Polyploidy promotes primary tumor growth, but opposes the evolution of Her2 independent growth

Our results thus far have focused on the role of polyploidy in the growth and survival of established recurrent tumors. We next sought to understand the impact of polyploidy on the development of recurrent tumors by focusing on the different stages of primary tumor growth, regression, and recurrence. We first assessed the impact of polyploidy on primary tumor growth. To address this, we digested primary MTB;TAN tumors and generated early-passage tumor cell cultures grown in the presence of dox to maintain Her2 expression. In light of recent work demonstrating increased chromosome mis-segregation during the proliferation of dissociated epithelial cells in culture [23], we first examined whether tumor cell ploidy changes during growth in vitro. We found that multiple tumor cell cultures derived from independent primary MTB;TAN tumors rapidly became dominated by tetraploid tumor cells (Supplemental Figure 4A and B), in spite of the fact that we never observed tetraploid cells in primary tumors in vivo (see Figure 1A).

We leveraged this this spontaneous polyploidization to generate matched cultures of diploid and tetraploid primary MTB;TAN tumor cells and compare their tumorigenic potential. Spontaneously formed tetraploids and diploids were sorted based on DNA content (Supplemental Figure 4C) and then injected orthotopically into the inguinal mammary gland of immunocompromised mice on doxycycline. The ploidy of the injected cell population was confirmed by DNA content analysis (Supplemental Figure 4D). Mice were palpated to monitor for tumor formation, and once tumors formed tumor volumes were measured twice per week using calipers. Tumors arising from the injection of tetraploid cells grew more quickly than tumors generated from diploid cell injections (Figure 6A and B; Welch’s t-test, p=0.0128). The proportion of Ki67 positive cells was significantly higher in WGD tumors (Supplemental Figure 4E; Welch’s t test, p = 0.0152). There were no significant differences in the proportion of apoptotic cells as indicated by immunohistochemical staining for cleaved caspase-3 (Supplemental Figure 4F; Welch’s t-test, p=0.06). These results suggest that while tetraploid tumor cells do not spontaneously occur in primary MTB;TAN tumors in vivo, when tetraploidy is induced experimentally, tetraploidy promotes the growth of primary MTB;TAN tumors.

**Figure 6.**
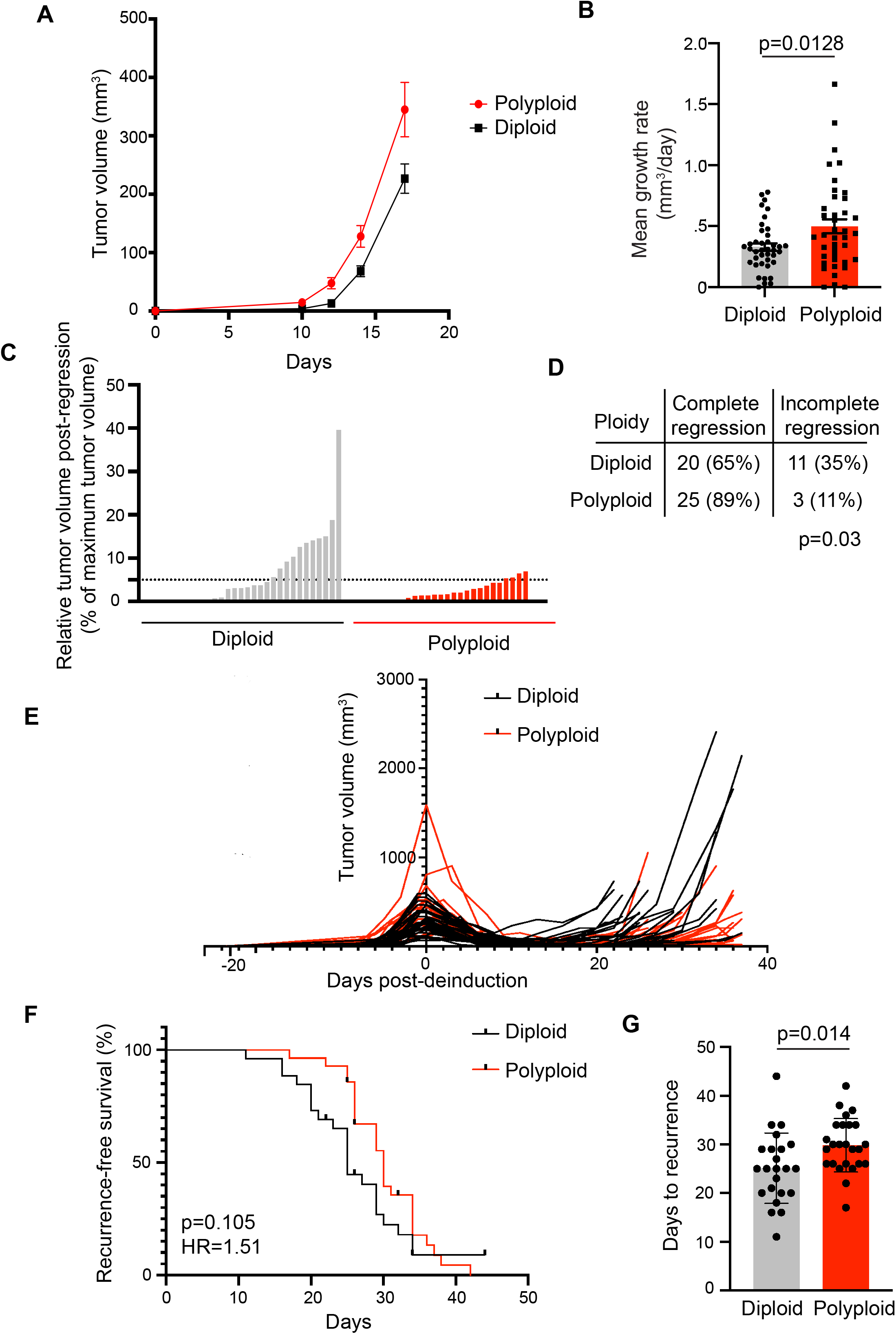
Polyploidy promotes primary tumor growth but opposes tumor recurrence. A) Tumor volume curves for primary tumors (Her2 on; +dox) in mice injected with diploid or tetraploid tumor cells. B) Mean growth rate of diploid and tetraploid primary tumor growth. Significance was determined using Welch’s t-test. C) Minimum tumor volume following tumor regression in diploid and tetraploid tumors. The extent of tumor regression was calculated by dividing the minimum tumor volume following Her2 downregulation by the maximum tumor size. The dotted line represents 95% tumor regression. D) The fraction of completely or incompletely regressing tumors in each cohort. Incomplete regression was defined as a minimum tumor volume > 5% of the tumor volume at de-induction. Differences in complete regression between cohorts as determined using Fisher’s exact test. E) Tumor volume curves for individual diploid and recurrent tumors. Mice were injected with cells between day −150 and −100 and were removed from doxycycline to de-induce Her2 at Day 0. F) Kaplan-Meier plot showing recurrence-free survival for diploid and tetraploid tumors following Her2 downregulation. Hazards Ratio (HR) and p-value were determined using a log-rank test. G) Median number of days to recurrence for diploid and tetraploid tumors. Significance was determined using Student’s t-test. *p<0.05

In light of our finding that approximately one-third of recurrent tumors have a substantial tetraploid cell population, we next asked how tetraploidy influences tumor recurrence. To do this, we removed doxycycline from the drinking water of mice with diploid or tetraploid primary tumors, leading to Her2 downregulation and mimicking treatment with anti-Her2 targeted therapy. Tumor volume was monitored over time. Significantly more diploid tumors failed to regress in response to downregulation of the Her2, presumably having evolved oncogene independent growth and survival mechanisms at a higher frequency than tetraploid tumors (Figure 6C and D; Fisher’s exact test, p=0.03). In tumors that did regress, tumor volume continued to be monitored and time to tumor recurrence was measured (Figure 6E). In contrast to the growth-promoting effects of tetraploidy on primary tumors, tetraploid tumors recurred at slightly later time points than diploid tumors (Figure 6F and G; log-rank test, p=0.105, HR=1.51; Student’s t-test comparing median recurrence-free survival times, p=0.014). Differences in proliferation and cell death between cohorts, as measured by Ki67 and cleaved caspase 3 staining, respectively, were non-significant (Supplemental Figure 4G and H; Welch’s t-test, p=0.039 for Ki67; p=0.3 for cleaved caspase-3).

### Polyploid tumor gene expression suggests altered immune interactions

To understand the mechanistic basis of the accelerated growth of tetraploid primary tumors, we first assessed relative levels of aneuploidy in diploid and tetraploid tumors. Metaphase spreads were performed on early-passage diploid and tetraploid primary tumors. Tetraploid tumors showed evidence of a higher degree of aneuploidy, likely indicative of increased chromosomal instability, consistent with our observations in vitro and with previous reports in the literature (Figure 7A).

**Figure 7.**
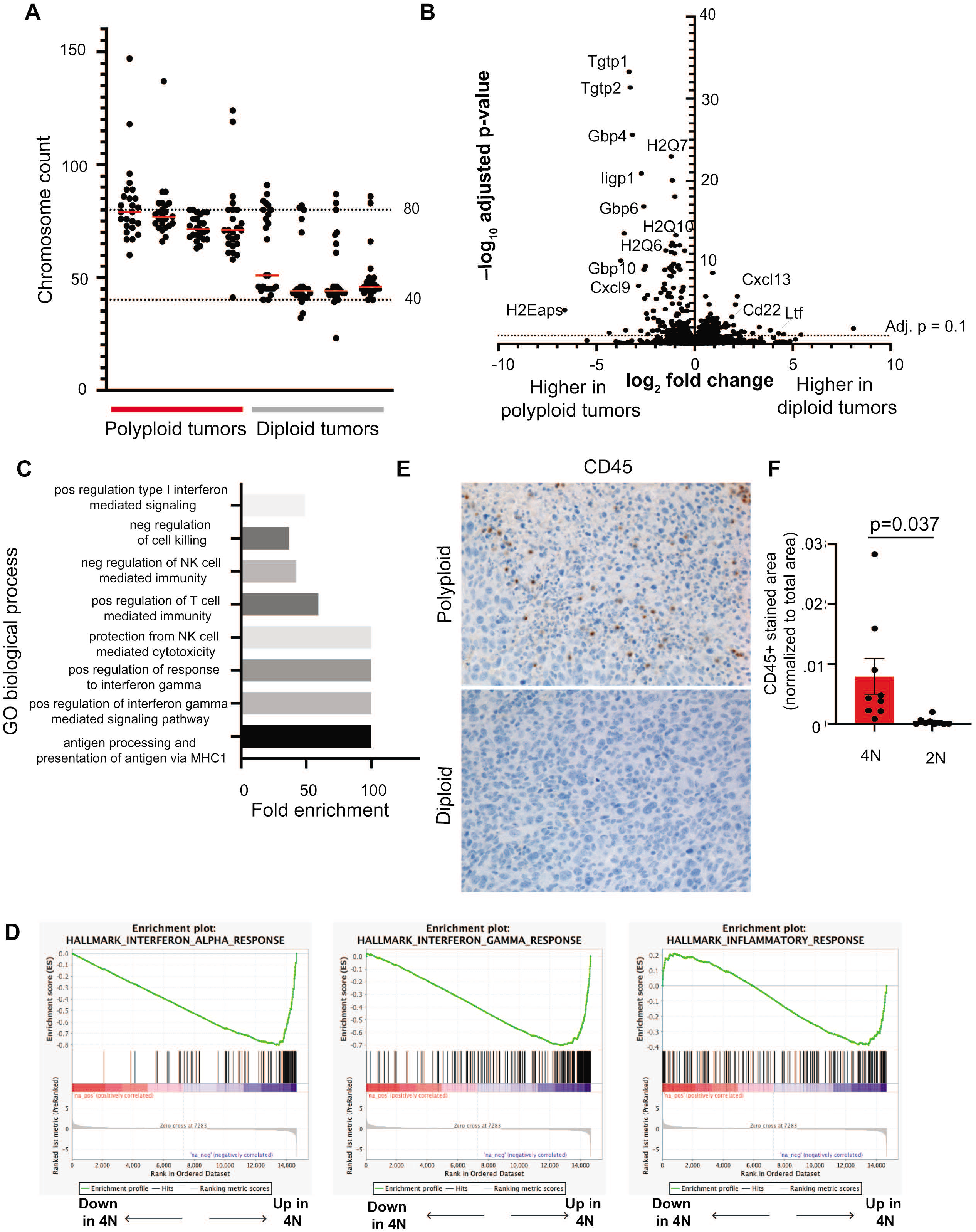
Polyploid primary tumors exhibit increased chromosomal instability and evidence of an innate immune response. A) Chromosome counts from metaphase spreads performed on early passage diploid and tetraploid primary tumors. Red bars represent median chromosome counts. Dashed lines represent diploid and tetraploid chromosome counts. B) Volcano plot showing differentially expressed genes between diploid and tetraploid tumors. Fold-change is shown on x-axis and adjusted p-value as determined by DESeq2 is shown on y-axis. The dotted line indicates an adjusted p-value of 0.1. C) Selected GO Biological Process terms from the top 50 significantly enriched in tetraploid tumors (FDR < 0.05). D) Selected gene sets enriched in tetraploid tumors by GSEA analysis (FDR q-value < 0.1). E) Representative images of CD45 staining on diploid and WGD primary tumors. F) Quantification of CD45 staining on diploid and WGD primary tumors. CD45 staining area is normalized to the total nuclear area. The mean and standard error of the mean are represented for each group. Significance was determined using Welch’s t-test.

The proliferation of tetraploid and highly aneuploid cells is opposed by numerous intrinsic and extrinsic tumor suppressive pathways. Thus, we hypothesized that tumors arising from polyploid cells would reflect selection for cells that are able to overcome these growth-suppressive mechanisms. To gain insight into pathways that permit the growth of polyploid tumors, we next performed RNA sequencing of diploid and polyploid primary and recurrent tumors. A number of genes were upregulated in primary polyploid tumors relative to primary diploid tumors (Figure 7B). No differentially expressed genes were found between diploid and polyploid recurrent tumors. Genes significantly upregulated in polyploid primary tumors included MHC class I genes, such as H2Q7, H2Q6 and H2Q10, and genes regulated by IFNg signaling, including guanylate GTPases Tgtp1, Tgtp2, Gbp4, Gbp6, and Gbp10 (Figure 7B). A smaller number of genes was downregulated in tetraploid tumors, including Cxcl13 and Cd22, which regulate B cell function, and Ltf, which is produced by neutrophils. GO term analysis of genes significantly upregulated in tetraploid tumors identified processes such as type I interferon signaling, negative regulation of natural killer mediated immunity, positive regulation of response to IFNγ, and antigen processing and presentation of antigen via MHC I (Figure 7C, Supplemental Table 1). Gene set enrichment analysis further demonstrated that tetraploid tumors are enriched in gene sets related to IFNα and IFNγ signaling, and a general immune response (Figure 7D and Supplemental Table 3). These results suggested that tetraploid tumors induce an immune response characterized by elevated IFNγ signaling and an associated upregulation of antigen presentation via MHC I. To directly assess immune cell infiltration into these tumors, we performed staining for the pan-leukocyte marker CD45. We observed a higher proportion of CD45+ staining in polyploid primary tumors, reflecting increased immune infiltration (Figure 7E and F; Welch’s t-test, p =0.037). Because athymic nude mice used as hosts for these experiments lack adaptive immunity, increased CD45 staining likely reflects the presence of innate immune cells. Together these results suggest that the accelerated growth of polyploid primary tumors is associated with increased aneuploidy and innate immune cell infiltration, consistent with other findings [11].

## Discussion

In the current study, using a genetically engineered mouse model of Her2-driven breast cancer and recurrence, we found that nearly 40% of recurrent tumors arising following Her2 downregulation undergo whole-genome duplications. In contrast, WGD was never observed in primary tumors in this model. To study the functional consequences of WGD in recurrent tumors, we created matched models of diploid and polyploid recurrent tumors. We found that near-tetraploid recurrent tumors exhibited increased chromosomal instability, decreased proliferation and increased survival in stress conditions. The effects of WGD on tumor growth were variable. Surprisingly, WGD slowed tumor growth in recurrent tumors, while primary WGD tumors grew more quickly. Our results highlight the context-dependent effects of WGD on tumor growth.

Our finding that WGD is associated with advanced disease, specifically recurrent tumors that form following Her2 downregulation, is consistent with reports from human cancers. Analysis of genomic sequencing data has shown that WGD is associated with increased mortality across cancer types [2]. WGD is detected at elevated levels in triple negative and Her2+ breast cancers [4], aggressive subtypes of breast cancer, and has been shown to play a role in driving resistance to chemotherapies and targeted therapies [24–26]. Despite the strong association of WGD with aggressive and advanced tumors, the basis for this relationship remains unknown. We propose that polyploid recurrent tumors that arise in MTB;TAN mice can provide a tractable model for deciphering the role of WGD in tumor progression.

Numerous studies have shown that WGD is associated with aneuploidy [6, 8, 9, 19], and that the proliferation of aneuploid cells resulting from WGD is opposed by growth suppressive mechanisms [12, 20, 27–29]. We made similar observations in our models. We found that polyploid recurrent tumor cells in both the spontaneous and induced models exhibit increased aneuploidy and increased rates of chromosome instability. Consistent with this, polyploid cells in both models had modest growth defects, both in vitro and in vivo. This suggests that aneuploidy and proliferation defects are two invariant consequences of WGD, whether it occurs spontaneously during tumor evolution or through experimental perturbations.

Our results suggest that the timing of WGD during tumor growth and recurrence is a critical determinant of its effect on tumor growth. Although we did not detect spontaneous WGD in primary tumors, the induction of polyploidy in primary tumors accelerated their growth. This suggests that the absence of WGD in primary tumors is because there is not a sufficient frequency of cell division errors that give rise to WGD, and not because primary tumor growth selects against WGD. Recurrent tumors that arise in this model undergo an epithelial-to-mesenchymal transition (EMT) [17]. Interestingly, a recent study found that sustained cell proliferation during EMT can lead to increases in mitotic errors [30]. It is possible that WGD in recurrent tumors may be a consequence of mitotic defects during the EMT process, which likely occurs following oncogene downregulation or in residual tumors [31].

Our observation that recurrent tumors frequently exhibit WGD, but that polyploid recurrent tumors grow more slowly than their diploid counterparts, was surprising. An optimal level of CIN may be necessary to promote tumor progression. High levels of chromosomal instability introduced by the experimental induction of WGD may exceed levels that would provide a fitness advantage and instead prove deleterious. Our data may be consistent with the occasionally contradictory relationship between CIN and patient outcomes. It has been previously observed that in ovarian cancer, gastric cancer, nonsmall cell lung cancer, and ER-breast cancer, patients with the highest levels of CIN actually had a better prognosis as compared to patients with intermediate CIN levels [32].

We speculate that aneuploidy and CIN may prove beneficial in certain conditions that cancer cells encounter during tumor progression, such as low nutrient or low growth factor signaling conditions. Consistent with this, we showed the WGD promotes the survival of cancer cells in response to low serum in vitro. The residual cancer cells that survive Her2 downregulation have decreased mitogenic and survival signaling [33], which may represent a cellular stress that selects for WGD. We propose a model where WGD is selected for during the dormant residual tumor stage yet slows the growth of established, proliferating tumors. Indeed, WGD has proven advantageous in adaptation to stress conditions in diverse model systems [21, 25, 34]. However, in the absence of such stressful conditions, WGD negatively affects cell fitness.

The net effect of WGD in tumor growth may also be dependent on the strength of oncogenic signaling in the cancer cell. In our model, primary tumors with high levels of Her2 signaling may be capable of overcoming the growth-suppressive effects of WGD, while recurrent tumors lacking Her2 signaling cannot. Increased growth factor signaling has previously been implicated as a mechanism to overcome G1 arrest resulting from cytokinesis failure [12, 35]. The effect of WGD may also depend on the evolutionary barriers that need to be overcome for tumor recurrence. Tumors that undergo WGD have increased chromosomal instability, but also extra copies of tumor suppressor genes. Extra copies of tumor suppressors have been shown to mediate the tumor suppressive effects of polyploidy in mouse models of liver cancer [36]. Previous work has demonstrated that loss of tumor suppressor genes is required for tumor recurrence following oncogene withdrawal [37, 38]. It is possible that polyploidy delays tumor recurrence by preventing the loss of these tumor suppressor genes.

Our RNA-seq data showed that polyploid tumors had activation of IFNα, IFNγ, and antigen presentation pathways, suggesting increased infiltration of immune cells in these tumors, which we confirmed through CD45 staining. These results are reminiscent of findings by other groups showing that polyploidy is associated with increased immunogenicity [27] and that aneuploidy leads to upregulation of NK cell ligands [11]. Interestingly, in our model this immune response was associated with accelerated rather than slowed tumor growth. This raises the possibility that polyploidy may promote primary tumor growth in the absence of an adaptive immune system, but slow primary tumor growth in immunocompetent mice, which would be consistent with previous findings [8, 27]. Indeed, recent studies examining human tumors have shown that established aneuploid tumors actually have fewer adaptive immune cells and diminished host immune responses [39, 40] suggesting that these tumors evolve immune escape mechanisms. Together these results suggest the consequences of polyploidy, including aneuploidy, shape interactions between tumors and the microenvironment, leading to changes in gene expression that are likely to influence tumor growth and characteristics.

Taken together, our study highlights the importance of identifying the context-dependent effects of WGD on tumor progression. Defining the cooperating genetic mutations and cellular environment that allow cells to overcome the barriers to polyploid tumor cell growth and proliferation is critical to understanding how WGD shapes tumor evolution. These insights may yield new strategies to target polyploidy, a frequent genetic alteration that is associated with poor prognosis in diverse cancers.

## Materials and Methods

### Mice

All experiments were approved by Duke IACUC (Approval #A199-17-08 and #A152-20-07) and performed in compliance with ARRIVE guidelines. Husbandry conditions were consistent with ranges recommended by The Guide for the Care and Use of Laboratory animals. Mice were kept on a 12-hour light and 12-hour dark schedule at 20-26° C and relative humidity of 40-70%. Female mice were used for all experiments at 6-7 weeks of age.

Primary and recurrent tumors in MMTV-rtTA;TetO-Her2/neu (MTB;TAN) mice were generated as previously described [17, 41, 42]. Briefly, MTB;TAN mice were administered doxycycline at 6 weeks of age. Primary tumors were harvested at volume of 500-3000 mm^3^. To generate recurrent tumors, mice with primary tumors were removed from dox to induce Her2 downregulation and tumor regression. Mice were palpated weekly to monitor for tumor recurrence, and recurrent tumors were harvested at a volume of approximately 100-4500 mm^3^.

### Orthotopic tumor growth and recurrence assays

For orthotopic tumor growth experiments, 500,000 diploid or near tetraploid cells were injected bilaterally into the inguinal mammary fat pad of nu/nu mice. Nu/nu mice were obtained from the Duke University Breeding Core. Mice were palpated weekly until the formation of a palpable tumor, then tumor dimension were measured using calipers a minimum of twice a week to monitor growth. Mice were sacrificed when tumors reached between 10 and 12 mm in diameter,. Tumor volumes were calculated using the formula ((π × length × width^2^)/6). Mean growth rates were calculated using the formula (AUC – (vol_1_ × day_n_))/(day_n_^2^).

For orthotopic recurrence assays, 500,000 diploid or tetraploid MTB;TAN primary tumor cells were injected bilaterally into the inguinal mammary fat pad of nu/nu mice. Doxycycline was added to the drinking water of mice two days prior to injection at a concentration of 2 mg/ml with 5% sucrose. The mice remained on dox through the course of primary tumor growth. Mice were palpated weekly until the formation of a palpable tumor, then a minimum of twice a week with calipers to monitor growth. When tumors reached approximately 10 millimeters by 10 millimeters in diameter, one cohort of diploid and near tetraploid primary tumors were sacrificed. Remaining cohorts had doxycycline removed from the drinking water, initiating Her2 downregulation. Tumors that regressed to a minimum size of less than 5% of the maximum tumor volume were considered regressed. Cohorts were monitored for the appearance of recurrent tumors and tumor growth was monitored as previously noted. Mice were sacrificed when recurrent tumors reached 10 millimeters by 10 millimeters in diameter. Primary and recurrent tumors were harvested and fixed in formalin for paraffin embedding and immunohistochemistry, digested to create cell lines and snap frozen for RNA isolation.

### DNA content analysis

For DNA content analysis, single cell suspensions were generated via enzymatic digestion [43]. Briefly, tumor tissue was minced and digested in collagenase and hyaluronidase (StemCell Technologies). After rinsing in media, cells were resuspended in red blood cell lysis buffer and incubated at room temperature. Cells were then rinsed in media then resuspended in DNAse (100 ug/mL) and Dispase II (5 mg/mL), and then resuspended in media and counted. Where noted, CD45 positive cells were depleted from tumor cell digests using the EasySep ^™^ Mouse streptavidin Rapidspheres^™^ Isolation Kit (Stem Cell Technologies, Catalog #19860) and a biotin conjugated anti Mouse CD45 antibody (Stem Cell Technologies, Catalog #60030BT) according to the manufacturers protocol.

Cells were resuspended in PBS at a concentration of 1 x 10^6^ cells per ml of PBS. Ice cold 100% ethanol was added, at a ratio of 3 ml of ethanol to 1 ml of PBS, dropwise while mixing. Cells were stored at −20°C for a minimum of 2 hours before analysis. Prior to staining, cells were rinsed twice in PBS and then resuspended in 1 ml DNA staining buffer (0.01% propidium iodide, 0.1% sodium citrate, 0.3% Triton-X 100, 2 mg/ml RNase A) per million cells. Cells were incubated at room temperature for 15 minutes in the dark. Samples were then analyzed on a FACS Canto A analyzer (BD Biosciences).

Data were analyzed using FCS Express software (DeNovo). Cell populations were gated to select for live cells (FSC-A x SSC-A) and single cells (FSC-H x FSC-A). The single-cell population was additionally refined by selecting for single cells based on PI-A x PI-W. DNA histograms were plotted for PI-A and the cell cycle distribution and presence of aneuploid/hyperdiploid cells estimated using FCS Express MultiCycle software (DeNovo). The distribution was autofitted to the optimal model.

To separate cells based on DNA content, cells were stained for 1 hour at 37° C with Vybrant DyeCycle Violet (ThermoFisher) at a concentration of 10 μM in the dark. Cells were filtered with a 40 micron strainer prior to sorting. Cells were excited with a UV laser on a BD DiVA cell sorter. Diploid cells (2N DNA content) and tetraploid cells (8N DNA content) were sorted after gating for live cells and single cells.

### Cell culture

Recurrent tumor cells were grown in DMEM with 10% SCS, 1% Pen-Strep, 1% glutamine and supplemented with EGF (0.01 ug/ml, Sigma) and insulin (5 ug/ml, Gemini Bioproducts). Primary tumor cells were grown in DMEM with 10% FBS, 1% Pen-Strep, 1% glutamine and supplemented with EGF (.01 ug/ml, Sigma), insulin (5 ug/ml), hydrocortisone (1 ug/mL), progesterone (1 uM), prolactin (5ug/ml) and doxycycline (2 ug/ml) to maintain Her2 expression.

For colony formation assays, cells were counted and suspended at a concentration of 100 cells per ml. 10 milliliters of each cell suspension were plated in 10-centimeter dishes in full serum media. The following day, cells were rinsed twice in PBS and the full serum media was replaced with DMEM with 10% SCS, 5% Pen Strep and 5% glutamine in one cohort, and reduced serum media with 5%, 2.5% or 1.25% SCS, 5% Pen Strep and 5% glutamine media. Cells were grown for 10 days and media was refreshed every 3 days. Plates were rinsed twice in PBS then stained for 10 minutes at room temperature with crystal violet. Plates were dried, scanned and colonies quantified using Fiji [44].

For cell counting experiments, 50,000 cells from the induced model or 100,000 cells from the spontaneous model were seeded in triplicate in six well plates. Cells were counted daily using a hemocytometer. Each cell population was measured three times.

### Fluorescent in situ hybridization (FISH)

Digested tumor cells were fixed in a 3:1 ratio mixture of methanol and glacial acetic acid and stored at −20° C. FISH was performed using Kreatech FISH probes (Leica Biosystems) for regions of chromosomes 2 and 11 (KI-30501) and for chromosomes X and 16 (KI-30503) for each tumor according to manufacturer’s protocol. Cells were stained with DAPI prior to the addition of mounting media. Slides were imaged using at a magnification of 100x and excited with 488, 405 and 561 lasers. The chromosome 16 specific mouse FISH probe was direct labeled with PlatinumBright^™^ 495. The chromosome 11 specific mouse FISH probe was direct labeled with Platinum Bright^™^ 550. A minimum of 25 cells were scored for each tumor for probes for chromosome 2 and 16.

### Metaphase spreads and karyotyping

For metaphase spreads, 150,000 cells were seeded in 6 well plates. The following day cells were treated with colchicine (Karyomax, ThermoFisher 15212012) at a concentration of 1 μg/ml and incubated for 1 hour at 37° C. Media was removed and cells were rinsed and trypsinized. Cell pellets were collected and resuspended in 5 ml of ice cold .56% potassium chloride solution, inverted and incubated at room temperature for 6 minutes. Cells were pelleted and all but 100 microliters the supernatant was removed. The cells were resuspended in the remaining solution. 5 ml of methanol glacial acetic acid fixative (3:1) was then added dropwise while mixing. Following fixation, cells were again pelleted and all but 100 ul of supernatant removed. A small volume of the resuspended solution was pipetted onto glass slides and dried at room temperature for a minimum of 1 hour. DAPI and mounting media were added to each slide. A coverslip was added and sealed. Slides were imaged using a fluorescent microscope at 40x magnification with a UV light source Any chromosome counts less than 15 were excluded from scoring as technical artifacts. For recurrent tumor cultures, 75 spreads were counted for each sample. For sorted cells, a minimum of 50 spreads were counted for the spontaneous model and 23 spreads for the induced model. For cells cultured from orthotopic tumor digests, 25 spreads were evaluated per sample.

G-band karyotyping of recurrent tumor cell cultures was performed by KaryoLogic Inc (Durham, NC). Cytogenetic analysis was performed on twenty-five G banded metaphase spreads of each mouse cell line.

### BrdU staining

For BrdU staining, 100,00 cells were seeded in six well plates. The following day media conditions were switched to high (DMEM with 10% SCS, 5% Pen Strep and 5% glutamine) or low serum conditions (DMEM with 5%, 2.5% or 1.25% SCS). After 24 hours, BrdU (BD Pharmigen, 51-7581KZ) was added at a concentration of 10 μM to the media for 20 minutes before harvesting cells. Cells were trypsinized then fixed in 70% ethanol and stored at −20° C for at least 30 minutes. Prior to analysis, cells were resuspended in 5 ml of PBS and rehydrated at 4 degrees for 1 hour. Cells were denatured (2M HCl, .5% Triton in PBS) for 2 hours at room temperature. Cells were resuspended in neutralization solution (0.1 M sodium tetraborate decahydrate) pH 8.5) for 30 min. Cells were then resuspended in antibody solution (Alexa Fluor 488 Mouse anti-BrdU, 558599, in 1% BSA, 0.5% Tween 20 in PBS) and incubated for 1 hour at room temperature in the dark. 1 ml of PBS was added, and the cells were spun down and resuspended in 1 ml of DNA staining buffer per 1 million cells and passed through a 40-micron filter. After 30 minutes of incubation at room temperature, samples were analyzed by on a FACS Canto (BD Biosciences) analyzer. FlowJo (BD BioSciences) was used for flow cytometry analysis. Each condition was treated and analyzed in duplicate.

### Immunohistochemistry

Tumor tissue was fixed in 10% neutral buffered formalin for 15 hours, rinsed twice in PBS then stored in 70% ethanol prior to embedding in paraffin. Immunohistochemistry was performed on 5-μm thick sections using antibodies against mouse cleaved-caspase 3 (Cell Signaling #9661S, 1:400) and Ki67 (Thermo RM-9106-S, 1:200) by the Duke Research Immunohistology Shared Resource (Durham, NC).

Quantification of IHC staining was performed using Fiji [44]. Colors were deconvoluted to separate positive staining and hematoxylin. For positive staining and hematoxylin channels, images were converted into 8-bit images and a threshold for positive staining was applied across all images. The number of particles of positive staining was quantified. This positive staining quantification was normalized to the corresponding hematoxylin measurement for each image. Three regions of interest were arbitrarily selected for imaging and quantification from each tumor section.

### RNA-sequencing and gene expression analysis

RNA was isolated from tumors using the RNAeasy kit (Qiagen). RNA was sequenced using the Illumina NovaSeq 6000 library and sequencing platform with 50 base paired end reads by the Duke GCB Sequencing and Genomic Technologies Shared Resource (Durham, NC). Quality control, trimming, and alignment and generation of counts from sequencing data was performed as previously described [14]. Differential gene expression analysis was performed using DESeq2 [45]. Genes with an adjusted p-value of less than 0.05 were used for Gene Ontology analysis using the Gene Ontology online resource [46] [47]. To perform Gene Set Enrichment Analysis, all genes were pre-ranked based on the magnitude of differential expression between diploid and near tetraploid tumors and analyzed using the desktop version of GSEA software (Broad Institute).

### Statistical Reporting

GraphPad Prism9 was used to perform statistical tests and to generate graphs. Differences in survival were evaluated using a log-rank Mantel Cox test. Differences in variance were evaluated using an F-test. Comparisons between two groups were tested using a 2-tailed Welch’s t-test or student’s t-test. Comparisons between multiple groups were evaluated using a two-way ANOVA and Sidak’s multiple comparisons test. Significant results are those considered with a p-value of less than 0.05.

