## Supplemental Figure for "Context-dependent effects of whole-genome duplication during mammary tumor recurrence"

#### Supplementary Information

#### Supplemental Figure 1

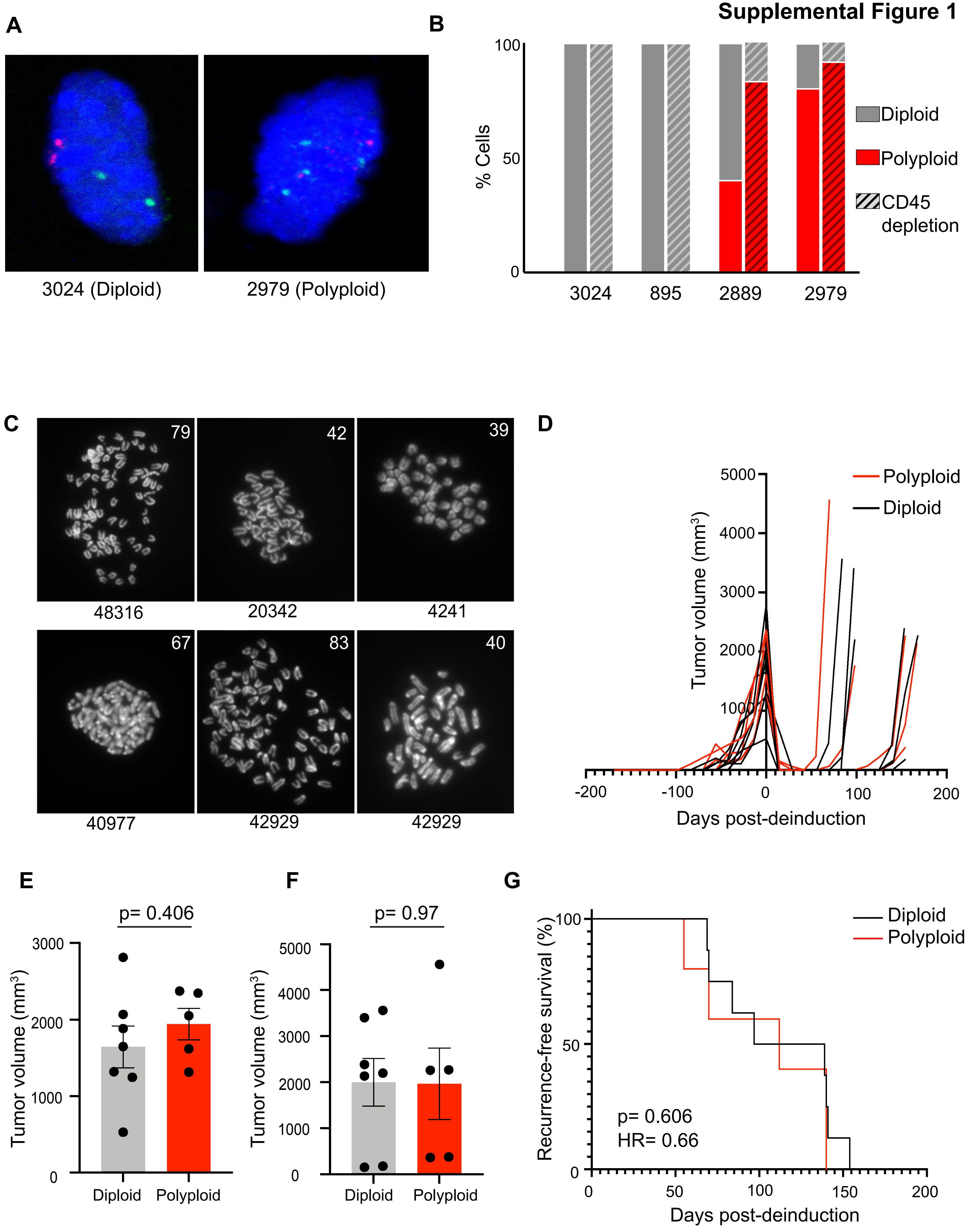

**Supplemental Figure 1. Tumor recurrence is associated with alterations in tumor cell ploidy.**
**A.** Representative FISH images for chromosome X (red) and chromosome 16 (green) on a diploid and tetraploid tumor. **B.** The frequency of diploid and tetraploid cells in tumors with or without depletion of CD45+ cells prior to DNA content analysis. **C.** Representative metaphase spreads of cells cultured from recurrent tumors arising in MTB;TAN mice. The number of chromosomes in each image is shown in the inset. **D.** Tumor volume curves for primary and recurrent tumors in MTB;TAN mice. Mice were administered doxycycline between day -150 and -100, and were removed from doxycycline to de-induce Her2 at Day 0. **E.** Tumor volume at de-induction of the antecedent primary tumor of recurrent tumors classified as diploid or tetraploid. Significance was determined using Welch's t-test. ($p=0.406$, Welch's t-test) **F.** Tumor volume at sacrifice of recurrent tumors classified as diploid or tetraploid ($p=0.97$, Welch's t-test). Significance was determined using Welch's t-test. **G.** Kaplan-Meier plot showing recurrence-free survival of diploid and polyploid recurrent tumors following Her2 de-induction. Hazards Ratio (HR) and p-value were determined using a log-rank test.

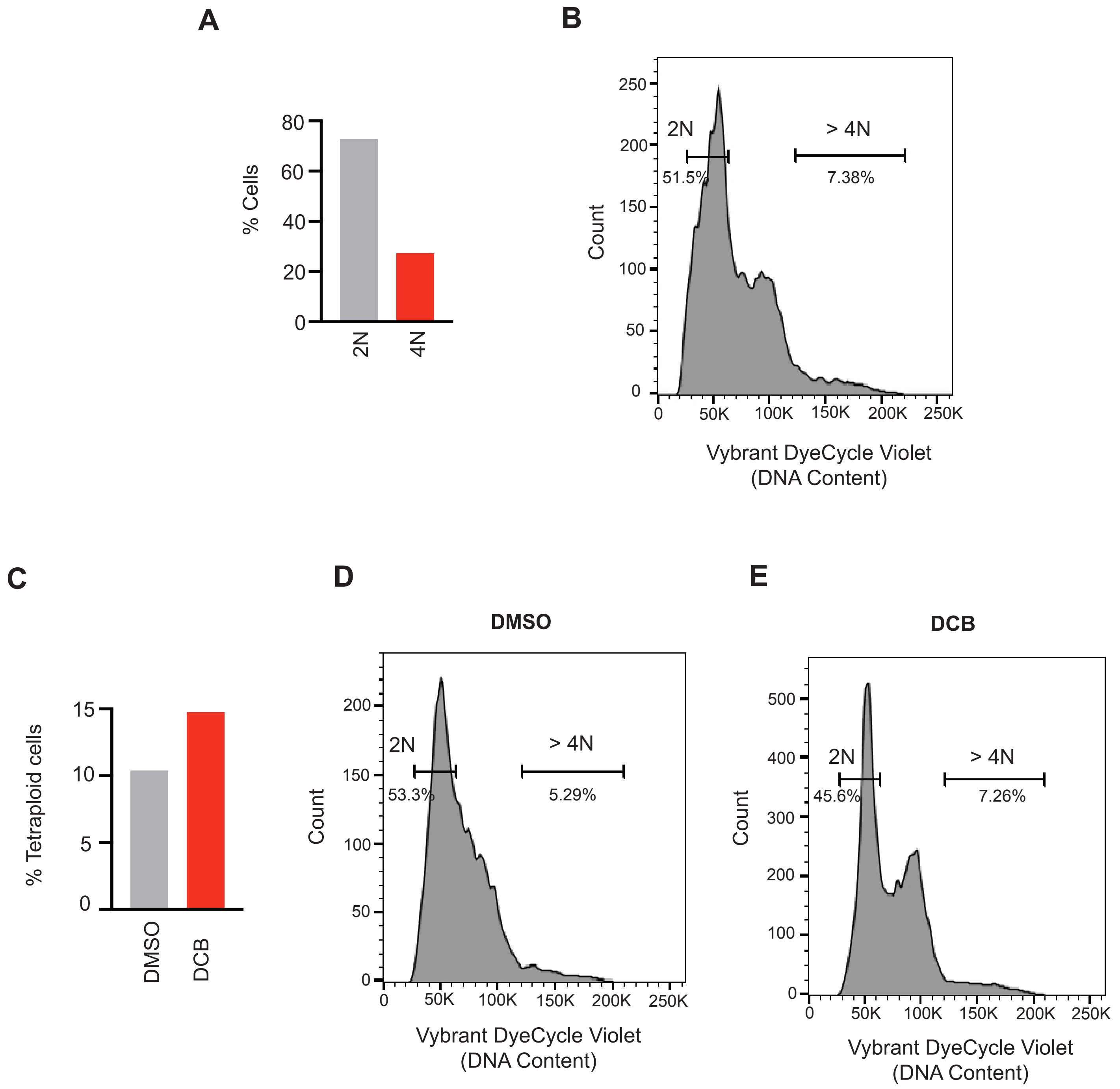


**Supplemental Figure 2. Creation of matched diploid and tetraploid recurrent tumor cell models.** **A.** The percentage of diploid and spontaneously formed tetraploid cells in unsorted 42929 cultures. DNA content analysis was performed, and cell cycle modeling used to estimate the percentage of diploid and tetraploid cells. **B.** Live-cell DNA content analysis of a mixed population of diploid and spontaneously formed near tetraploid (or WGD) cells. **C.** The percentage of diploid and induced tetraploid cells in unsorted diploid cells treated with DMSO or DCB. Quantification of DNA content analysis. DNA content analysis was performed, and cell cycle modeling used to estimate the percent of diploid and tetraploid cells. **D.** DNA content analysis of diploid cells treated with DMSO. **E.** DNA content analysis of diploid cells treated with DCB for 16 hours at 1 μM.

### Supplemental Figure 3

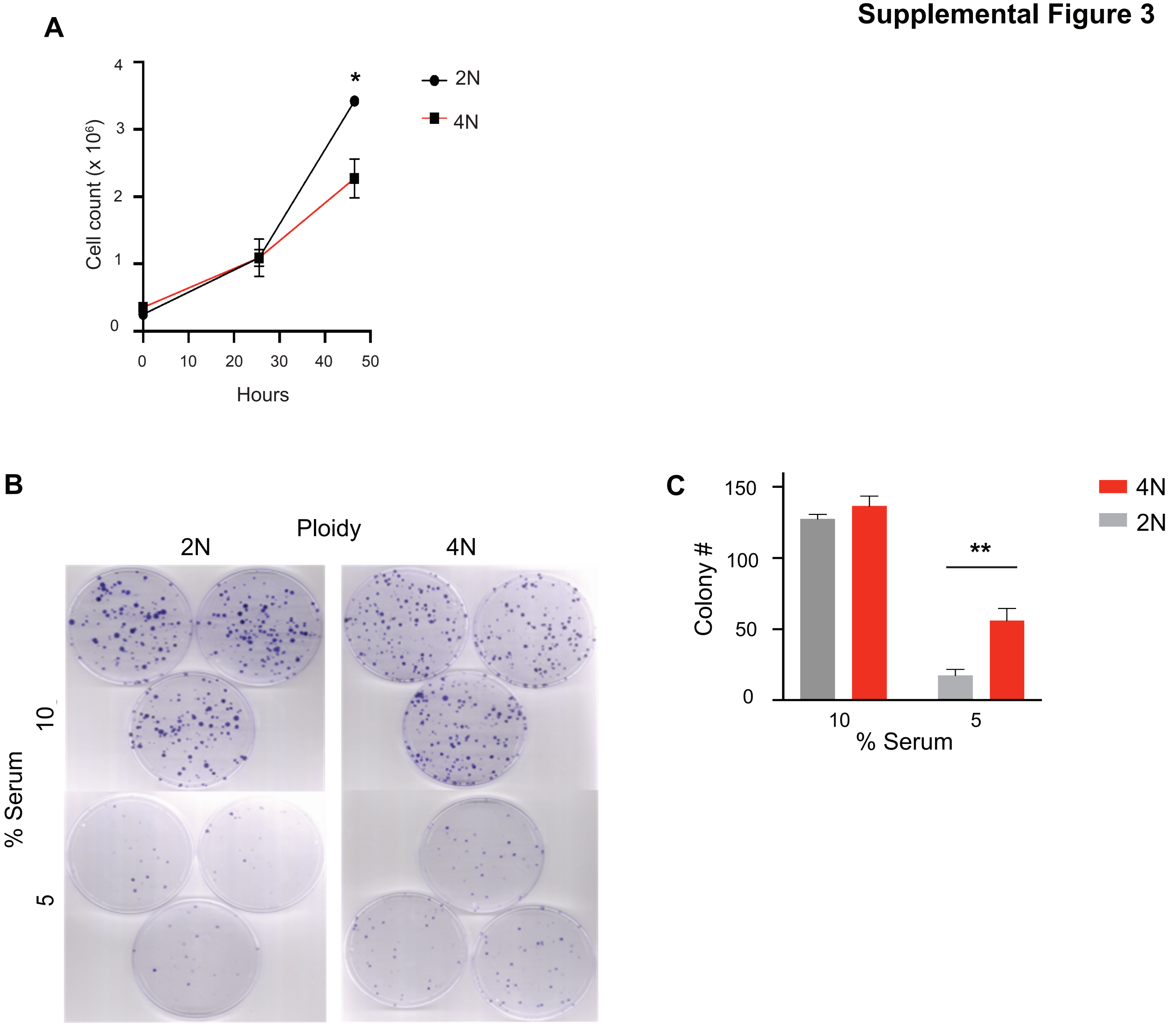


**Supplemental Figure 3. Polyploid recurrent tumor cells demonstrate a competitive advantage in low serum culture.**
**A.** Cell count of diploid and spontaneously formed tetraploids over time. Significance was determined using two-way ANOVA with Sidak's multiple comparison test. **B.** Colony formation assays of diploid and induced tetraploid cells in various serum concentrations. **C.** Quantification of colony number from B. Significance was determined using Student's t-test.

*$p<0.05$, **$p<0.01$

### Supplemental Figure 4

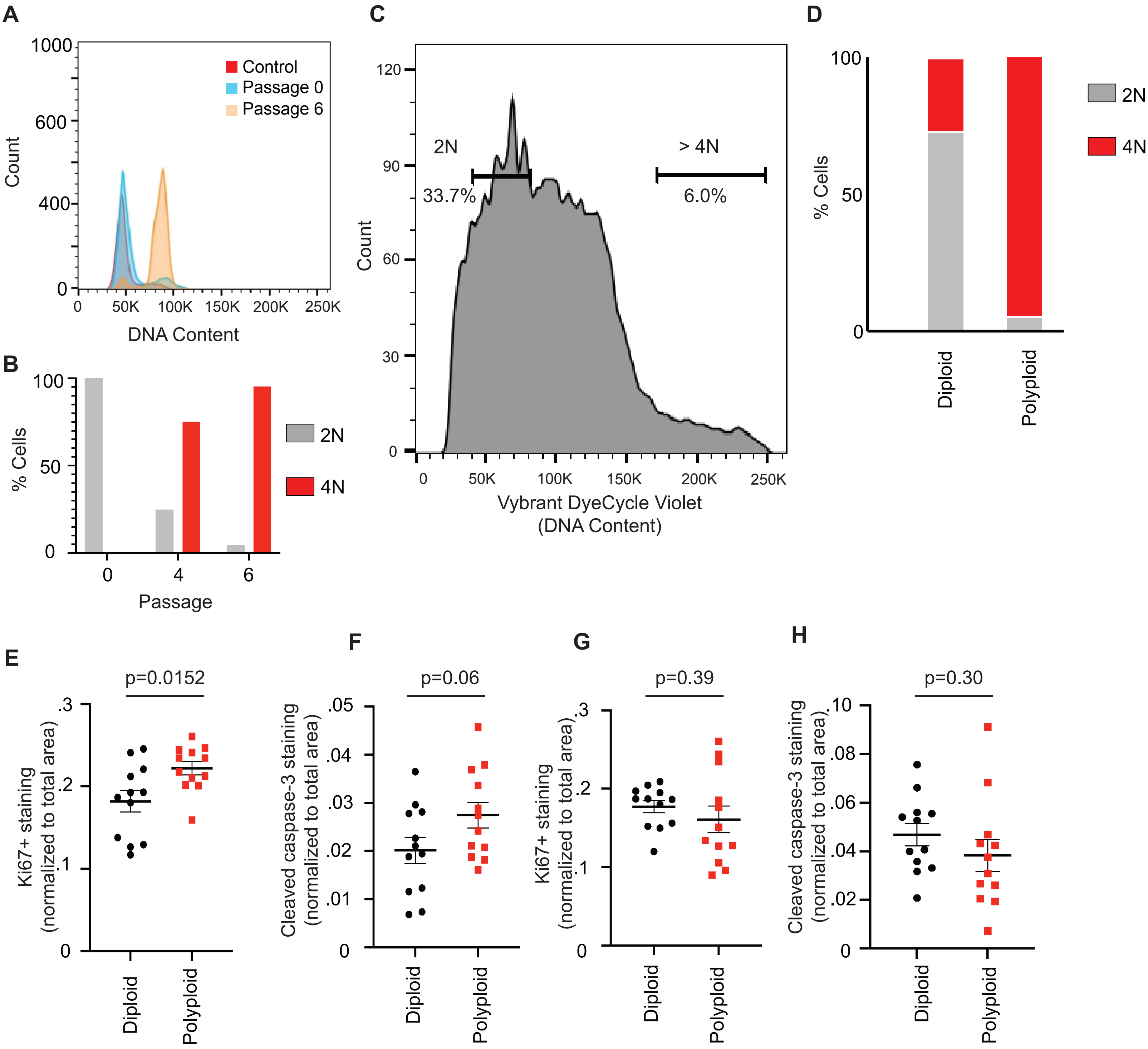


**Supplemental Figure 4. Polyploidy promotes primary tumor growth but opposes tumor recurrence.**
**A.** DNA content analysis of primary tumor cells at early passages compared to a near-diploid control. **B.** Quantification of the percentage of diploid and WGD cells in populations of primary tumor cells at various passages. Cell cycle modeling was used to quantify data from DNA content analysis. **C.** Live-cell DNA content analysis of a mixed population of diploid and spontaneously formed near tetraploid (or WGD) cells. Gates indicated cells sorted for injection. **D.** Quantification of the percentage of diploid and WGD cells in populations of injected primary tumor cells. Cell cycle modeling was used to quantify data from DNA content analysis. **E.** The proportion of area positive for Ki67 staining in diploid and WGD primary tumors. **F.** The proportion of area positive for cleaved caspase 3 staining in diploid and WGD primary tumors. **G.** The proportion of area positive for Ki67 staining in diploid and WGD recurrent tumors. **H.** The proportion of area positive for cleaved caspase 3 staining in diploid and WGD recurrent tumors.

For E-H, significance was determined using Student's t-test.

#### Supplemental Table 1. Gene Ontology enrichment analysis showing the top 50 gene ontology terms enriched in WGD tumors.

| GO biological process complete | Fold Enrichment | FDR |
|---|---|---|
| antigen processing and presentation of endogenous peptide antigen via MHC class I via ER pathway, TAP-dependent (GO:0002485) | > 100 | 3.50E-04 |
| antigen processing and presentation of exogenous protein antigen via MHC class Ib, TAP-dependent (GO:0002481) | > 100 | 3.46E-04 |
| antigen processing and presentation of endogenous peptide antigen via MHC class Ib via ER pathway, TAP-dependent (GO:0002489) | > 100 | 1.72E-02 |
| antigen processing and presentation of endogenous peptide antigen via MHC class Ib via ER pathway (GO:0002488) | > 100 | 1.70E-02 |
| antigen processing and presentation of exogenous peptide antigen via MHC class I, TAP-dependent (GO:0002479) | > 100 | 1.69E-02 |
| antigen processing and presentation of exogenous peptide antigen via MHC class Ib (GO:0002477) | > 100 | 6.85E-06 |
| regulation of antigen processing and presentation of peptide antigen via MHC class I (GO:0002589) | > 100 | 2.65E-02 |
| positive regulation of interferon-gamma-mediated signaling pathway (GO:0060335) | > 100 | 3.03E-05 |
| positive regulation of response to interferon-gamma (GO:0060332) | > 100 | 2.99E-05 |
| protection from natural killer cell mediated cytotoxicity (GO:0042270) | > 100 | 2.95E-05 |
| MHC protein complex assembly (GO:0002396) | > 100 | 3.91E-02 |
| cytosol to endoplasmic reticulum transport (GO:0046967) | > 100 | 3.88E-02 |
| antigen processing and presentation of peptide antigen via MHC class Ib (GO:0002428) | 94.24 | 2.40E-22 |
| antigen processing and presentation of endogenous peptide antigen (GO:0002483) | 93.09 | 4.62E-25 |
| antigen processing and presentation of endogenous peptide antigen via MHC class I (GO:0019885) | 93.09 | 3.85E-25 |
| antigen processing and presentation of endogenous peptide antigen via MHC class I via ER pathway (GO:0002484) | 90.88 | 8.11E-21 |
| antigen processing and presentation of endogenous peptide antigen via MHC class Ib (GO:0002476) | 90.88 | 7.63E-21 |
| antigen processing and presentation of endogenous peptide antigen via MHC class I via ER pathway, TAP-independent (GO:0002486) | 87.31 | 2.98E-19 |
| antigen processing and presentation of endogenous antigen (GO:0019883) | 86.75 | 9.32E-25 |
| antigen processing and presentation via MHC class Ib (GO:0002475) | 84.82 | 8.63E-22 |
| antigen processing and presentation of peptide antigen via MHC class I (GO:0002474) | 79 | 2.84E-25 |
| adhesion of symbiont to host (GO:0044406) | 75.73 | 5.09E-06 |
| antigen processing and presentation of exogenous peptide antigen via MHC class I (GO:0042590) | 70.68 | 3.20E-03 |
| cellular response to interferon-beta (GO:0035458) | 67.67 | 2.33E-19 |
| positive regulation of T cell mediated cytotoxicity (GO:0001916) | 65.25 | 1.78E-20 |
| defense response to protozoan (GO:0042832) | 63.04 | 1.02E-13 |
| response to interferon-beta (GO:0035456) | 61.1 | 2.16E-21 |
| regulation of T cell mediated cytotoxicity (GO:0001914) | 59.52 | 5.37E-20 |
| antigen processing and presentation of peptide antigen (GO:0048002) | 58.39 | 1.24E-23 |
| response to protozoan (GO:0001562) | 56.89 | 2.52E-13 |
| regulation of interferon-gamma-mediated signaling pathway (GO:0060334) | 56.55 | 3.06E-04 |
| regulation of response to interferon-gamma (GO:0060330) | 56.55 | 3.02E-04 |
| regulation of ribonuclease activity (GO:0060700) | 56.55 | 2.99E-04 |
| antigen processing and presentation of exogenous peptide antigen (GO:0002478) | 54.98 | 5.07E-08 |
| antigen processing and presentation of exogenous antigen (GO:0019884) | 49.89 | 4.90E-09 |
| positive regulation of type I interferon-mediated signaling pathway (GO:0060340) | 48.93 | 7.74E-03 |
| positive regulation of T cell mediated immunity (GO:0002711) | 45.85 | 1.79E-18 |
| negative regulation of natural killer cell mediated cytotoxicity (GO:0045953) | 44.64 | 6.38E-04 |
| antigen processing and presentation (GO:0019882) | 44.43 | 3.79E-25 |
| negative regulation of natural killer cell mediated immunity (GO:0002716) | 42.41 | 7.54E-04 |
| positive regulation of leukocyte mediated cytotoxicity (GO:0001912) | 41.38 | 7.10E-18 |
| negative regulation of leukocyte mediated cytotoxicity (GO:0001911) | 36.88 | 1.21E-03 |
| positive regulation of cell killing (GO:0031343) | 35.71 | 5.40E-17 |
| regulation of T cell mediated immunity (GO:0002709) | 34.62 | 8.15E-17 |
| regulation of leukocyte mediated cytotoxicity (GO:0001910) | 34.33 | 7.32E-18 |
| negative regulation of leukocyte mediated cytotoxicity (GO:0001911) | 36.88 | 1.21E-03 |
| positive regulation of cell killing (GO:0031343) | 35.71 | 5.40E-17 |
| regulation of T cell mediated immunity (GO:0002709) | 34.62 | 8.15E-17 |
| regulation of leukocyte mediated cytotoxicity (GO:0001910) | 34.33 | 7.32E-18 |
| response to type I interferon (GO:0034340) | 33.93 | 1.61E-03 |
| type I interferon signaling pathway (GO:0060337) | 33.48 | 1.87E-02 |
| cellular response to type I interferon (GO:0071357) | 33.48 | 1.85E-02 |
| regulation of nuclease activity (GO:0032069) | 31.41 | 2.05E-03 |
| negative regulation of cell killing (GO:0031342) | 31.41 | 2.03E-03 |

**Supplemental Table 2. Gene Ontology enrichment analysis showing all gene ontology terms enriched in non-WGD tumors.**

| GO biological process complete | Fold enrichment | FDR |
|---|---|---|
| cell adhesion (GO:0007155) | 4.44 | 5.46E-02 |
| biological adhesion (GO:0022610) | 4.39 | 4.12E-02 |
| cellular component organization (GO:0016043) | 2.03 | 7.82E-02 |
| cellular component organization or biogenesis (GO:0071840) | 1.95 | 4.99E-02 |

**Supplemental Table 3. Hallmark gene sets enriched in tetraploid primary tumors (FDR q value < 0.1).**

| Name | Nominal Enrichment Score | Nominal p-value | FDR q-value |
|---|---|---|---|
| HALLMARK_INTERFERON_ALPHA_RESPONSE | -2.7800088 | 0 | 0 |
| HALLMARK_INTERFERON_GAMMA_RESPONSE | -2.6772268 | 0 | 0 |
| HALLMARK_MYC_TARGETS_V1 | -2.1526678 | 0 | 0 |
| HALLMARK_OXIDATIVE_PHOSPHORYLATION | -2.018973 | 0 | 5.56E-04 |
| HALLMARK_MYOGENESIS | -1.7201589 | 0 | 0.008518 |
| HALLMARK_ALLOGRAFT_REJECTION | -1.7123371 | 0 | 0.00758 |
| HALLMARK_MYC_TARGETS_V2 | -1.687307 | 0.008492569 | 0.009487 |
| HALLMARK_DNA_REPAIR | -1.625411 | 0.002320186 | 0.015754 |
| HALLMARK_IL6_JAK_STAT3_SIGNALING | -1.6135163 | 0.00443459 | 0.015725 |
| HALLMARK_KRAS_SIGNALING_DN | -1.5830822 | 0.004210526 | 0.018791 |
| HALLMARK_INFLAMMATORY_RESPONSE | -1.4571732 | 0.011135858 | 0.052256 |
